# Activation of hypoactive parvalbumin-positive fast-spiking interneuron restores dentate inhibition to prevent epileptiform activity in the mouse intrahippocampal kainate model of temporal lobe epilepsy

**DOI:** 10.1101/2024.04.05.588316

**Authors:** Sang-Hun Lee, Young-Jin Kang, Bret N. Smith

**Affiliations:** Department of Biomedical Sciences, Colorado State University, Fort Collins, CO 80523, USA

**Keywords:** temporal lobe epilepsy, GABAergic interneuron, parvalbumin-expressing basket cell, dentate gyrus

## Abstract

Parvalbumin-positive (PV+) GABAergic interneurons in the dentate gyrus provide powerful perisomatic inhibition of dentate granule cells (DGCs) to prevent overexcitation and maintain the stability of dentate gyrus circuits. Most dentate PV+ interneurons survive status epilepticus, but surviving PV+ interneuron mediated inhibition is compromised in the dentate gyrus shortly after status epilepticus, contributing to epileptogenesis in temporal lobe epilepsy. It is uncertain whether the impaired activity of dentate PV+ interneurons recovers at later times or if it continues for months following status epilepticus. The development of compensatory modifications related to PV+ interneuron circuits in the months following status epilepticus is unknown, although reduced dentate GABAergic inhibition persists long after status epilepticus. We employed PV immunostaining and whole-cell patch-clamp recordings from dentate PV+ interneurons and DGCs in slices from male and female sham controls and intrahippocampal kainate (IHK) treated mice that developed spontaneous seizures months after status epilepticus to study epilepsy-associated changes in dentate PV+ interneuron circuits. We found that the number of dentate PV+ cells was reduced in IHK treated mice. Electrical recordings showed that: 1) Action potential firing rates of dentate PV+ interneurons were reduced in IHK treated mice up to four months after status epilepticus; 2) Spontaneous inhibitory postsynaptic currents (sIPSCs) in DGCs exhibited reduced frequency but increased amplitude in IHK treated mice; and 3) The amplitude of evoked IPSCs in DGCs by optogenetic activation of dentate PV+ cells was upregulated without changes in short-term plasticity. Video-EEG recordings revealed that IHK treated mice showed spontaneous epileptiform activity in the dentate gyrus and that chemogenetic activation of PV+ interneurons abolished the epileptiform activity. Our results suggest not only that the compensatory changes in PV+ interneuron circuits develop after IHK treatment, but also that increased PV+ interneuron mediated inhibition in the dentate gyrus may compensate for cell loss and reduced intrinsic excitability of dentate PV+ interneurons to stop seizures in temporal lobe epilepsy.

**Highlights:**

- Reduced number of dentate PV+ interneurons in TLE mice
- Persistently reduced action potential firing rates of dentate PV+ interneurons in TLE mice
- Enhanced amplitude but decreased frequency of spontaneous IPSCs in the dentate gyrus in TLE mice
- Increased amplitude of evoked IPSCs mediated by dentate PV+ interneurons in TLE mice
- Chemogenetic activation of PV+ interneurons prevents epileptiform activity in TLE mice

## 1. Introduction

In the dentate gyrus, powerful parvalbumin-positive (PV+) interneuron mediated perisomatic inhibition is essential for preserving sparse firing of dentate granule cells (DGCs) under normal circumstances (Espinoza et al., 2018; Hu et al., 2014). Dentate PV+ interneurons are classified into two subtypes: PV+ basket cells (PVBCs) and axo-axonic cells (AACs) (Houser, 2007). These cells form symmetrical synapses on the somata and proximal dendrites of DGCs and interneurons and the axon initial segments of DGCs, respectively (Freund and Buzsaki, 1996; Pelkey et al., 2017). Dentate PV+ interneuron mediated perisomatic inhibition limits DGC excitability under physiological conditions (Lee et al., 2019), but interneuron loss and compromised activity of surviving dentate PV+ interneurons contribute to disinhibition of DGCs and have been critically implicated in spontaneous seizure expression in temporal lobe epilepsy (TLE) (Kobayashi and Buckmaster, 2003; Marx et al., 2013; Proddutur et al., 2023; Zhang and Buckmaster, 2009). Other research suggests that there are fewer AAC axon terminals in TLE (Alhourani et al., 2020; Ribak, 1985), in agreement with loss of PV+ interneurons in TLE. Conversely, neuroanatomical studies suggest that dentate PV+ interneurons form increased symmetrical synapses on axon initial segments of DGCs, maintain a similar number of synapses on DGC somata, and undergo axon sprouting in TLE (Christenson Wick et al., 2017; Thind et al., 2010; Wittner et al., 2001). It remains unknown whether and how plasticity in surviving dentate PV+ interneurons functionally compensates for cell loss and compromised activity in this interneuron population.

Activation of hippocampal PV+ interneurons prevents ongoing seizure activity in TLE (Krook-Magnuson et al., 2013; Wang et al., 2018). Thus, gaining a deeper understanding of the function of dentate PV+ interneurons in TLE is highly significant. Findings from epileptic tissues suggest that hippocampal PV+ interneurons may remain hypoactive due to downregulated intrinsic and/or synaptic excitability in TLE. The "dormant dentate basket cell" hypothesis postulates that PVBCs lose excitatory inputs and become inactive or dormant in TLE (Sloviter, 1987; Sloviter, 1991), but there is disagreement on this hypothesis (Bernard et al., 1998). More recent studies showed that dentate PV+ interneurons (e.g., AACs) underwent reduced action potential (AP) generation along with reduced synaptic excitation early after experimental status epilepticus (Proddutur et al., 2023). However, it is uncertain whether dentate PV+ interneurons recover normal fast spiking weeks after status epilepticus (i.e., during the chronic phase of TLE). Dentate PV+ interneurons not only undergo reduced synaptic excitation, but also participate in downregulated PV+ interneuron mediated inhibition of DGCs early after status epilepticus and in epileptic tissues (Proddutur et al., 2023; Zhang and Buckmaster, 2009). However, surviving dentate PV+ interneurons manifest reduced AP generation, but increased synaptic excitation mediated by DGCs weeks after epileptogenic insults (e.g., traumatic brain injury), thus contributing to increased feedback inhibition in injured mice (Kang et al., 2022). It is unclear whether there are similar dentate PV+ interneuron associated compensatory changes in the chronic phase of TLE not associated with a traumatic brain injury.

In the present work, we used opto- and chemo-genetic methods in the intrahippocampal kainate (IHK) mouse model of TLE to determine: 1) the epilepsy-associated changes in overall dentate PV+ interneuron mediated inhibition in the context of PV+ interneuron loss in TLE and 2) the impact of activating PV+ interneurons on epileptiform activity in the dentate gyrus. Our opto- and chemo-genetic strategies targeting PV+ interneurons do not distinguish between PVBCs and AACs. We thus combined all dentate PV+ interneurons for the data analysis to determine if there were epilepsy-associated alterations in intrinsic excitability and synaptic excitation of dentate PV+ interneurons using whole-cell patch-clamp recordings.

## 2. Materials and methods

### 2.1. Animals

All procedures involving mice used in the current study were approved by the Institutional Animal Care and Use Committee of Colorado State University. All mice were maintained at ambient temperature and humidity with a 12-hour light/12-hour dark cycle and fed standard chow *ad libitum*. Three bi-transgenic mouse lines were employed in this work to target PV+ cells for optogenetics, chemogenetics, and patch-clamp recordings. We crossed a homozygous PVCre strain (stock #017320, The Jackson Laboratory) to a homozygous Ai32 mouse strain (expressing a ChR2/EYFP fusion protein in a Cre recombinase-dependent manner; stock #024109, The Jackson Laboratory), a homozygous G_q_-DREADD strain (expressing a G_q_- DREADD/mCitrine in a Cre recombinase-dependent manner; stock #024109, The Jackson Laboratory), or a homozygous red reporter strain (expressing a tdTomato in a Cre recombinase-dependent manner; stock #007909, The Jackson Laboratory). In this study, the resulting bi-transgenic offsprings are referred to as “PVCre::ChR2”, “PVCre::G_q_-DREADD”, and “PVCre::tdTomato”, respectively. C57BL/6J mice (stock #000664, The Jackson Laboratory) were used as negative controls for chemogenetic experiments. Similar numbers of males and females were used for this study, thus reducing sex bias (Will et al., 2017). The combined data from both sexes are presented in the current study. Reporting of *in vivo* Experiments (ARRIVE) guidelines and the Basel declaration, including the 3R (Replacement, Reduction, Refinement) concept, were considered when planning the experiments.

### 2.2. IHK mouse model of TLE and video monitoring for behavioral seizures

Seven-week-old male and female mice were anesthetized under isoflurane and unilaterally injected with 100–150 nl of sterile 0.9% saline or kainate (20 mM in 0.9% saline; Tocris Biosciences) into the dorsal hippocampus using the following stereotaxic coordinates relative to the bregma (anterior-posterior, –2.0 mm; medial-lateral, –1.25 mm; dorsal-ventral, –1.60 mm) as previously described (Kang et al., 2021). In order to minimize loss of mice caused by status epilepticus (>30 minutes of Racine stage 3–5 seizure), diazepam (7.5 mg/kg) was injected intraperitoneally 1 hour after the first Racine stage 5 seizure, characterized by tonic-clonic convulsions, rearing, running, jumping, and falling (Racine, 1972). Saline-injected mice (i.e., sham controls) also received intraperitoneal injections of diazepam. All mice were individually housed after surgery and recovered in the vivarium. Three to four weeks after kainate injection, spontaneous behavioral activities were video recorded continuously for 3 days (72 hr) using video cameras and Sirenia software (version 1.7.10; Pinnacle Technology) without implanting EEG electrodes. IHK mice that displayed Racine stage 3–5 seizures at least once during the 3-day video recordings were defined as “TLE mice” and used in this study.

### 2.3. Quantification of PV+ interneuron somata in the dentate gyrus

Since we performed whole-cell patch-clamp recordings using a restricted area of the ventral hippocampus (dorsal-ventral: approximately –2.4 to –3.6 mm from Bregma), we performed quantification of PV+ interneuron number using a similar area of the ventral hippocampus from 6 saline (3 females and 3 males) and 7 IHK (3 females and 4 males) injected mice. Horizontal slices from the ventral hippocampus (300 μm thickness) were placed in fixative solution (4% paraformaldehyde, 0.2% picric acid in 0.1 M phosphate buffer, pH 7.4) and stored at 4°C for 24– 48 hours. After washing in phosphate buffer, fixed brain slices were embedded in OCT Compound (Fisher Scientific) and resectioned at 30μm thickness using a cryostat (HM 550, Microm). The sections were then subjected to immunohistochemistry using primary (goat anti-PV, PVG213, 1:2000, Swant, Burgdorf) and secondary (donkey anti-goat alexa fluor 488, A11055, 1:500, Thermo Fisher Scientific) antibodies.

Tissue sections were blocked in 5% normal donkey serum in Tris-buffered saline (pH 7.4) containing 0.3% Triton-X 100 for 1 hour at room temperature, followed by incubations in primary and secondary antibodies diluted in Tris-buffered saline with 0.3% Triton-X 100 and 1% normal donkey serum. After washing thoroughly in Tris-buffered saline with 0.05% Tween-20, the sections were mounted in Vectashield medium and subjected to epifluorescence image acquisition using an Olympus IX73 microscope. Using a 20× objective (UPlanFL N, NA 0.5; Olympus) controlled by a motorized microscope stage (TANGO 3 DESKTOP, Märzhäuser Wetzlar), tile-scanned images of the dentate gyrus were acquired using cellSens Dimension software (version 2.3, Olympus). PV+ interneurons from sham controls and TLE mice were manually counted in a double-blinded manner. The area of the dentate gyrus used for soma quantification was measured and the number of counted neurons was normalized to tissue area. Images were analyzed with ImageJ (version 1.54f) software.

### 2.4. Whole-cell patch-clamp recordings

Horizontal hippocampal slices (300 µm) from ventral hippocampi were prepared from both sexes of PVCre::ChR2, PVCre::G_q_-DREADD, and PVCre::tdTomato mice 6-8 weeks or 4 months after IHK or saline injection. Brain slices were cut in ice-cold (2-4°C) sucrose-containing, oxygenated (95% O_2_/5% CO_2_) artificial cerebrospinal fluid (ACSF), incubated for one hour at 33°C, and then stored at room temperature until electrical recording. ACSF contained, in mM, 85 NaCl, 75 sucrose, 25 glucose, 24 NaHCO_3_, 4 MgCl_2_, 2.5 KCl, 1.25 NaH_2_PO_4_, and 0.5 CaCl_2_. Hippocampal slices were transferred to a recording chamber in ACSF containing, in mM, 124 NaCl, 26 NaHCO_3_, 11 glucose, 3 KCl, 2 CaCl_2_, 1.3 MgCl_2_, and 1.4 NaH_2_PO_4_. Slices were visualized with an upright microscope (BX51WI, Olympus) with infrared-differential interference contrast optics. For optogenetic experiments, light (470 nm) was delivered through the epifluorescence port of BX51WI, using LED source (X-Cite Xylis, Excelitas Technologies).

To examine intrinsic and synaptic properties of PV+ interneurons in the dentate gyrus, whole-cell patch-clamp recordings from PV+ interneurons in the dentate gyrus were performed in current- or voltage-clamp configuration using a recording pipette solution containing (in mM) 126 K-gluconate, 4 KCl, 10 HEPES, 4 ATP-Mg, 0.3 GTP-Na, 10 phosphocreatine, and 5.4 biocytin with a pH of 7.2 and osmolarity of 290–295 mOsm, which in this study is referred to as pipette solution #1. Patch pipettes had resistances of 3–5 MΩ. All electrical recordings were made at 33°C using a MultiClamp 700B amplifier (Molecular Devices). Electrical signals were filtered at 3 or 10 kHz using a Bessel filter and digitized at 10 or 20 kHz with a Digidata 1440A analog-digital interface (Molecular Devices). Series resistance was carefully monitored; electrical recordings were discarded if the series resistance changed >20% or reached 25MΩ. The recorded traces were analyzed using Clampfit software (version 10.7.0, Molecular Devices) and Mini Analysis (version 6.0.7, Synaptosoft Inc.).

Intrinsic and synaptic properties of dentate PV+ interneurons were measured as we previously described (Kang et al., 2022). Briefly, their intrinsic properties were measured from voltage responses to a series of 1 second hyperpolarizing and depolarizing current steps (-200 to +600 pA, +40 pA increments) from –60mV. The following properties were examined using Clampfit software or Mini Analysis: (1) Resting membrane potential was measured from the average voltage after an equilibration period; (2) Input resistance was measured from voltage responses to small current steps (–40 to +40 pA in 40 pA increments. The slope of current-voltage curve was calculated at steady state (0.8–1.0 second from the start of current steps); (3) AP threshold, amplitude, and half-width were measured from the first three APs evoked by a 1 second depolarizing current step (+400 pA). AP threshold was defined as the point at which the derivative of the membrane potential (dV/dt) first exceeded its mean by two standard deviations during the interspike interval. AP amplitude was measured from threshold to peak; half-width was defined as the AP duration at one half AP amplitude; and (4) Frequency and amplitude of spontaneous excitatory postsynaptic currents (sEPSCs) in dentate PV+ cells (V_hold_ = –70 mV) were measured for 2 minutes using Mini Analysis. sEPSCs were manually detected using Mini Analysis and included for this study only if amplitudes of the synaptic currents were greater than three times root mean square noise level, as we previously described (Kang et al., 2022).

To examine PV+ interneuron mediated inhibition in the dentate gyrus, we employed PVCre::ChR2 mouse line for optogenetic activation of PV+ interneurons. Whole-cell patch-clamp recordings from DGCs that were localized within the outer one-half of the granule cell layer were performed in voltage-clamp configuration. Optogenetic experiments were conducted using the pipette solution #1. Blue light (2 ms duration) was delivered through a 40× objective to the dentate gyrus and hilus (light power density, 9.9 mW/mm^2^) as we previously described (Kang et al., 2022). The power density was estimated from the field of view (662 μm diameter) of 40× objective (LUMPlanFL N, NA 0.8; Olympus). Blue light was centered on the somata of recorded cells. Evoked inhibitory postsynaptic currents (eIPSCs) were measured at 0mV (i.e., near the reversal potential of ionotropic glutamate receptor-mediated events) (Kang et al., 2022). Ten light stimuli (2ms duration, 10s interval) were used to analyze eIPSC amplitude. To examine short-term plasticity of eIPSCs, trains of theta-frequency light stimuli (i.e., 20 light pulses at 10 Hz) were employed. In addition, whole-cell patch-clamp recordings from dentate PV+ cells were performed in current-clamp configuration to examine whether optogenetic activation of dentate PV+ cells produces fewer APs in TLE mice. APs in dentate PV+ cells were evoked from –60 mV by 10 light stimuli (2 ms duration, 10s interval).

Frequency and amplitude of spontaneous IPSCs (sIPSCs) were measured as we previously described (Kang et al., 2021). DGCs were recorded in voltage clamp (V_hold_ = –70 mV). The DGC pipette solution contained (mM) 90 K-gluconate, 40 CsCl, 10 phosphocreatine, 10 HEPES, 3.5 KCl, 2 Mg-ATP, 1.8 NaCl, 1.7 MgCl_2_, 0.4 Na_2_GTP, and 0.05 EGTA, with a pH 7.2, and osmolarity of 290mOsm. The reversal potential for Cl^-^ in this solution was calculated as -26 mV and sIPSCs were inward under this recording condition (Kang et al., 2021; Lee et al., 2014). Kynurenic acid (1 mM; Sigma-Aldrich) was added to the ACSF to block sEPSCs. Amplitude and frequency of sIPSCs in DGCs were measured using Mini Analysis. A 2 minute control recording before bath application of clozapine N-oxide (CNO, 5 μM) and a 2 minute recording in CNO for each DGC were used for measuring amplitude and frequency of sIPSCs.

### 2.5. Dentate PV+ interneuron identification

All dentate PV+ interneurons were first selected based on expression of tdTomato (in PVCre::tdTomato mice) and EYFP (in PVCre::ChR2 mice) using epifluorescence optics (Olympus) and verified by their fast-spiking firing pattern (Hu et al., 2014). Under infrared-differential interference contrast, their somata were situated close to the granule cell layer. Specifically, 22 dentate PV+ cells and 26 dentate PV+ cells are recorded from sham control and TLE mice, respectively. The somata of most dentate PV+ cells (Sham controls: 16 PV+ cells; TLE mice: 21 PV+ cells) were located at the border of inner molecular layer/granule cell layer or within the inner molecular layer. The somata of the remaining PV+ cells (Sham controls: 6 PV+ cells; TLE mice: 5 PV+ cells) were located within granule cell layer or at the border of granule cell layer/hilus. The location of dentate PV+ cells used for slice electrophysiology in the current study was thus similar between sham control and TLE mice. After electrical recordings with pipettes containing biocytin, the slices were fixed in a solution containing 4% paraformaldehyde and 0.2% picric acid in 0.1 M phosphate buffer (pH 7.4) and stored at 4°C for 24–48 hours. The slices were washed in phosphate buffer. Biocytin-filled dentate PV+ cells were stained with 1:500 diluted streptavidin conjugated with Alexa 488 or 594 (Invitrogen) and examined for neuronal morphology using an epifluorescence upright microscope (BX43, Olympus) equipped with UPLanFL N 40× objective lens (NA 0.75, Olympus).

In a subset of the studies, stained brain slices were washed thoroughly in phosphate buffer and followed by a passive clearing procedure to image biocytin-filled PV+ cells. Brain slices were cleared by incubating in the Delipidation Buffer (LifeCanvas Technologies, Cambridge) for 1–2 hours at room temperature. After rinsing with phosphate buffer, brain slices were incubated in 50% and 100% of EasyIndex solution (LifeCanvas Technologies) for 30 minutes, placed on a concave microscope slide and coversliped with 100% of EasyIndex solution. Z-stack images (319.5 µm × 319.5 µm with 2 µm step intervals) were acquired under a 20× objective (Plan-Apochromat, NA 0.8) using a Zeiss LSM800 confocal microscope (Carl Zeiss). Using ImageJ, z-stack images were projected into two dimensions and merged to assess the morphology of biocytin-filled PV+ interneurons (**Fig. 2A**).

Electrophysiological data in this study were obtained only from dentate PV+ cells that manifested preferential localization of their axon terminals in the granule cell layer. These dentate PV+ interneurons manifested basket-like axon terminals in the granule cell layer (Fig. 2A). No dentate PV+ cells displayed prominent vertical aggregations of axonal varicosities, a crucial morphological characteristic of AACs (Proddutur et al., 2023; Somogyi and Klausberger, 2005). It appears to be impossible to unequivocally distinguish PVBCs from AACs without using electron microscopy to identify PV+ interneuron synaptic targets (Somogyi and Klausberger, 2005). However, based on their location at the border of inner molecular layer/granule cell layer or within the inner molecular layer, some of these cells are most likely AACs (Proddutur et al., 2023). Therefore, it is likely that dentate PV+ cells used in this study were PVBCs and AACs as previously described (Proddutur et al., 2023).

### 2.6. Video-EEG recordings

Four months after IHK injection, mice were anesthetized with isoflurane and ipsilaterally implanted with a 2-channel depth electrode (Plastics One) in stereotaxic coordinates corresponding to the dorsal dentate gyrus (anterior-posterior –2.5 mm, medial-lateral +1.75 mm, dorsal-ventral – 2.0 mm with respect to bregma). Implanted electrodes were firmly secured on the skull using mounting screws and dental cement. One week after the surgery, the mice were monitored by synchronized video-EEG recording for 24 hours a day for 4 days. Electrodes were connected to the 3-pin head mount (Plastics One), which was connected to the customized Mouse Preamplifier (Pinnacle Technology). EEG data were acquired at a 500 Hz sampling frequency with 50 Hz low pass filter and 0.5 Hz high pass filter using 8401-HS Data Conditioning and Acquisition System (Pinnacle Technology). LED cameras were projected from the top of cages for video recordings.

Synchronized video-EEG files were acquired with Sirenia Acquisition software (version 2.2.7) and 8400 Tethered Mouse System (Pinnacle Technology). EEG data sets were exported using Sirenia Seizure Pro (Pinnacle Technology) as an EDF file to measure epileptiform activity using NeuroScore’s Seizure Module (version 3.4.0, DSI). Epileptiform activity is defined as a spontaneous increase in voltage (> the mean + 3× root mean square of a baseline period for 10 sec) with spiking lasting no less than 5 sec and no less than 1 Hz in frequency. Interictal periods were set to a minimum of 3 s.

Intraperitoneal injections of saline or CNO (1 mg/kg in a 0.9% saline solution) were given on the second or third day between 10 a.m. and noon. Number and cumulative duration of epileptiform activity for the 3 hours before and after saline or CNO administration were quantified to determine whether chemogenetic activation of PV+ cells altered epileptiform activity in the dentate gyrus of TLE mice.

### 2.7. Statistical analysis and scientific rigor

Paired or unpaired (as appropriate) two-tailed Student’s *t* tests were used. In cases in which the data did not show normal distribution based on the Shapiro–Wilk test, then the Wilcoxon’s signed-rank or Mann–Whitney *U* tests for paired and unpaired data, respectively, were used. Otherwise, two-way ANOVAs were followed by Tukey tests for mean comparisons. Data are presented as mean ± SEM. A p-value less than 0.05 was considered significant (\**p* < 0.05; \*\**p* < 0.01; \*\*\**p* < 0.001; ns, not significant). Statistical analyses were performed using Prism 9 (GraphPad Software).

To avoid bias and increase rigor, the investigators performing experiments were blind to the animal treatment and drug treatments were randomized as we previously described (Kang et al., 2021; Kang et al., 2022). Similar numbers of replicates per mouse were collected for whole-cell patch-clamp recordings to avoid potential biases toward the results from mice with more replicates being obtained.

## 3. Results

### 3.1. Quantification of dentate PV+ interneuron number of TLE mice

We showed that the unilateral dorsal IHK mouse model produced localized hippocampal sclerosis that was limited to ipsilateral dorsal hippocampus, without involving the ventral or contralateral hippocampus (Kang et al., 2021). However, ventral hippocampi manifest reduced overall GABAergic inhibition and loss of cholecystokinin-expressing basket cells, a major subtype of hippocampal GABAergic interneurons, in the TLE mice (Kang et al., 2021). These results suggest that the ventral hippocampus, remote from the sclerotic dorsal hippocampus, also undergoes neuronal circuit reorganization. In this study, we examined whether there was loss of PV+ interneurons in the ventral dentate gyrus using the unilateral dorsal IHK mouse model, since vulnerability of PV+ interneurons in the ventral dentate gyrus is unknown in this model.

We employed PV immunostaining. As previously reported (Buckmaster and Dudek, 1997; Houser, 2007), most PV+ somata in sham control mice were found at or close to the granule cell layer (**Fig. 1A**). The position of PV+ somata in TLE mice was similar to that of sham control mice (**Fig. 1B**). However, significantly fewer PV+ somata were seen in the ventral dentate gyrus of TLE mice compared to sham controls (**Fig. 1C**; Sham controls: 16.95±1.23/1 mm^2^, 43 sections from 6 mice; TLE: 13.57±0.79/1 mm^2^, 51 sections from 7 mice; *p*=0.0205). In the TLE mice, the results from ipsi- and contra-dentate gyrus were combined since the number of PV+ somata in the dentate gyrus was similar between ipsi- and contra-lateral dentate gyrus (Ipsilateral: 13.6±1.09/1 mm^2^, 28 sections from 7 mice; Contralateral: 13.53±1.18/1 mm^2^, 23 sections from 7 mice; *p*=0.9668). The present results suggest that there is loss of PV+ interneurons in the ventral dentate gyrus after unilateral dorsal IHK injection. Our current findings are generally consistent with previous research showing that the dentate gyrus has PV+ interneuron loss in areas that are ventral to the IHK injection site (i.e., intermediate locations between dorsal and ventral dentate gyrus) (Marx et al., 2013). Our current research and the 2013 Marx et al. findings, however, differ in a few significant ways. The previous studies found no significant differences in the number of PV+ somata in the ventral dentate gyrus, whereas our present studies showed that there was a significant reduction in PV+ somata in the ventral dentate gyrus. The 2013 Marx et al. used IHK-injected mice (whose behavioral seizures were not established) at 21 days following IHK injection. In contrast, the current investigations used IHK-injected mice showing spontaneous behavioral seizures 4 to 6 weeks after IHK injection, which were confirmed to have developed TLE. These findings suggest that spontaneous seizures develop at later times following IHK-induced status epilepticus and are associated with the loss of PV+ interneurons.

**Figure 1.**
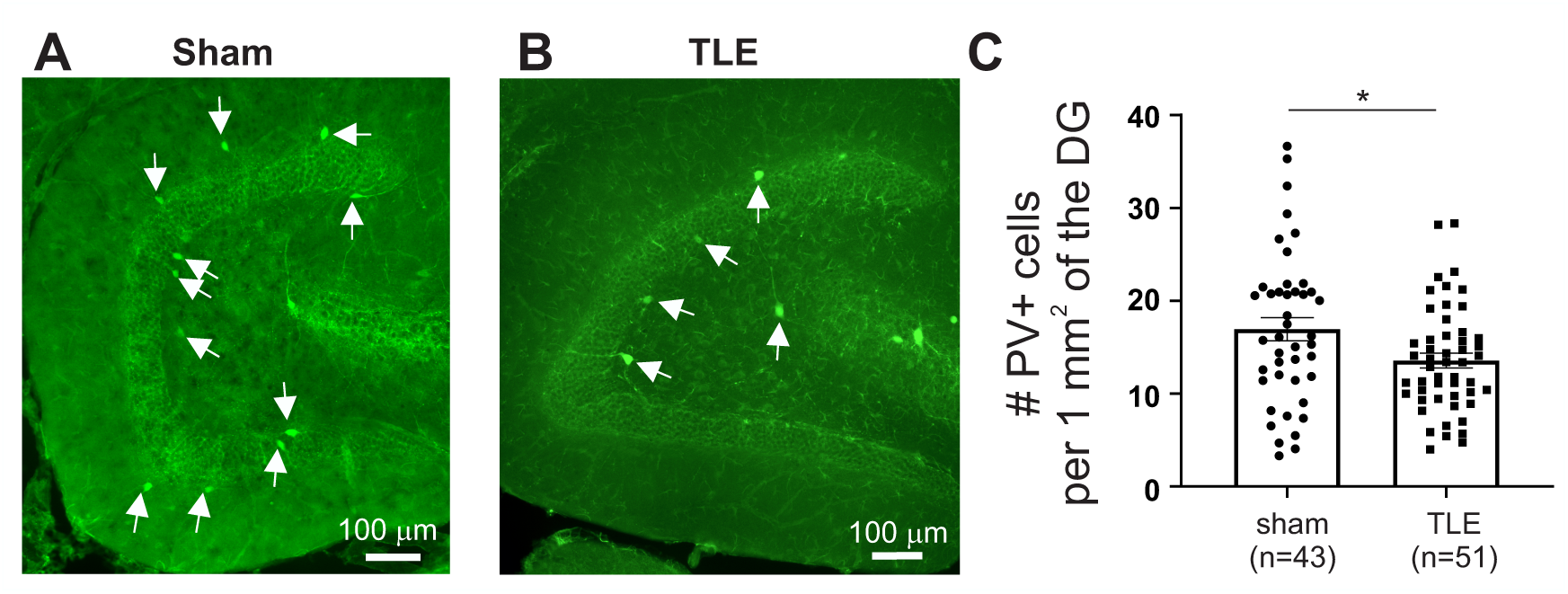
Reduced number of parvalbumin positive (PV+) interneurons in the dentate gyrus after intrahippocampal kainate (IHK) treatment. **(A, B)** PV+ interneurons were identified based on immunopositivity for PV (green). Representative images of the dentate gyrus from a sham control (Sham) or TLE mouse (TLE). In the sham control and TLE mice, most PV+ somata (marked with arrows) were found at or close to the granule cell layer. The number of PV+ neurons was significantly reduced in TLE mice. **(C)** Summary of the relative density of PV+ neurons in the dentate gyrus in sham control and TLE mice. The ipsi- and contralateral data were combined for each group. Solid circles and squares represent individual sections. The number of sections used for these studies was indicated by the numbers inside each parenthesis. Mean±SEM; \**p*<0.05.

### 3.2. Intrinsic excitability of surviving dentate PV+ interneurons of TLE mice

We performed whole-cell patch-clamp recordings from PV+ interneurons in the dentate gyrus of sham control and TLE mice 4-6 weeks after intrahippocampal saline or kainate injection to determine whether there were major changes in intrinsic properties of dentate PV+ interneurons in TLE. Dentate PV+ interneurons from sham control mice manifested high-frequency APs during current injection (**Fig. 2**). In sharp contrast, dentate PV+ interneurons ipsi- and contra-lateral to IHK injection manifested reduced frequency of evoked APs compared to sham controls (**Fig. 2**; two-way ANOVA with repeated measures; significant IHK treatment main factor, *p*=0.0019; significant IHK treatment×injected current amplitude interaction, *p*<0.0001; Sham controls: n=13 from 6 mice; TLE: n=14 from 6 mice). The results from ipsi- and contra-lateral dentate PV+ interneurons were pooled since the data from dentate PV+ interneurons ipsilateral to IHK injection did not differ from those from dentate PV+ interneurons contralateral to IHK injection (two-way ANOVA with repeated measurements; IHK treatment main factor, *p*=0.857). The frequency rate of APs induced by 400 pA depolarizing current steps was 41.9±5.7 Hz (n=14) in dentate PV+ interneurons from TLE mice, a substantial decrease compared to 66.1±2.3 Hz (n=13; *p*=0.003) in sham control mice. We also investigated whether other AP characteristics of dentate PV+ interneurons underwent significant changes. In TLE mice, there were significant increases in rheobase (Sham controls: 170.0±14.9 pA; n=13 from 6 mice; TLE: 225.7±21.2 pA; n=14 from 6 mice; *p*=0.048) and AP threshold (Sham controls: -40.7±0.7 mV; n=13 from 6 mice; TLE: - 37.7±0.7 mV; n=14 from 6 mice; *p*=0.011), whereas there were no significant alterations in other intrinsic membrane properties of dentate PV+ interneurons (**Table 1**). These findings demonstrated that lower firing frequency of surviving dentate PV+ interneurons was associated with increased rheobase and AP threshold in TLE mice.

**Figure 2.**
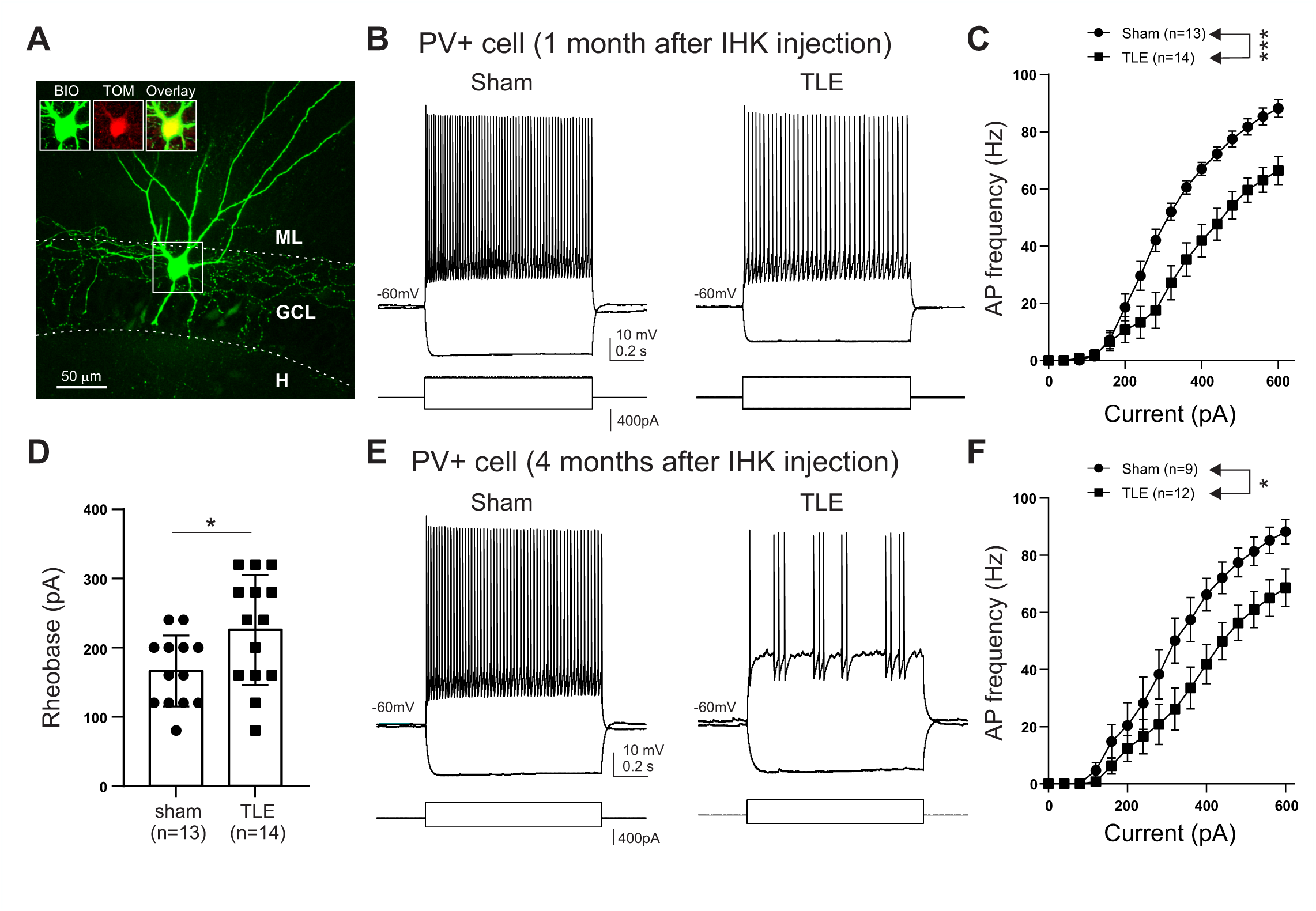
Reduced action potential (AP) frequency of dentate PV+ interneurons from TLE mice. **(A)** A representative image of a PV+ interneuron that was filled with biocytin (BIO) during a whole-cell patch-clamp recording in a transgenic mouse that expressed tdTomato (TOM) in PV+ interneurons. **(B)** One month following intrahippocampal saline and kainate injection, respectively, the voltage responses of dentate PV+ interneurons from sham control and TLE mice to depolarizing and hyperpolarizing current steps. (**C**) Summary of AP frequency of PV+ interneurons from recordings in (B). **(D)** Summary of rheobase of PV+ interneurons one month after IHK injection. Solid circles and squares represent individual PV+ interneurons. **(E,F)** AP firing properties of dentate PV+ interneurons of sham control and TLE mice 4 months after intrahippocampal saline and kainate injection, respectively (E). Summary of AP frequency of PV+ interneurons (F). Mean±SEM; \**p*<0.05; \*\*\**p*<0.005. Abbreviations: BIO, biocytin; GCL, granule cell layer; H, hilus; ML, molecular layer; TOM, tdTomato.

**Table 1.** Intrinsic properties of PV+ interneurons in sham controls and TLE mice666.

| 1 month after IHK or saline injection | Sham (n=13) | TLE (n=14) | Significant differences |
| --- | --- | --- | --- |
| RMP (mV) | -66.2 ± 2.2 | -67.4 ± 1.5 | $p=0.936$ |
| R<sub>input</sub> (MΩ) | 109.6 ± 6.1 | 99.9 ± 15.1 | $p=0.060$ |
| Rheobase (pA) | 170.0 ± 14.9 | 225.7 ± 21.22 | $p=0.048$ |
| C<sub>m</sub> (pF) | 111.1 ± 6.3 | 137.7 ± 12.9 | $p=0.080$ |
| AP threshold (mV) | -40.7 ± 0.7 | -37.7 ± 0.7 | $p=0.011$ |
| AP amplitude (mV) | 68.7 ± 2.0 | 66.5 ± 1.0 | $p=0.311$ |
| AP half-width (ms) | 0.59 ± 0.03 | 0.57 ± 0.04 | $p=0.711$ |
| AP frequency (Hz) | 66.1 ± 2.3 | 41.9 ± 5.7 | $p=0.003$ |
| 4 months after IHK or saline injection | Sham (n=9) | TLE (n=12) | Significant differences |
| RMP (mV) | -67.2 ± 1.2 | -63.2 ± 2.0 | $p=0.148$ |
| R<sub>input</sub> (MΩ) | 124.2 ± 12.1 | 107.2 ± 8.9 | $p=0.263$ |
| Rheobase (pA) | 191.1 ± 28.1 | 240.0 ± 25.1 | $p=0.165$ |
| C<sub>m</sub> (pF) | 112.8 ± 10.3 | 150.6 ± 18.1 | $p=0.218$ |
| AP threshold (mV) | -37.6 ± 1.1 | -37.9 ± 1.8 | $p=0.651$ |
| AP amplitude (mV) | 60.9 ± 2.3 | 64.4 ± 3.1 | $p=0.424$ |
| AP half-width (ms) | 0.62 ± 0.03 | 0.59 ± 0.03 | $p=0.385$ |
| AP frequency (Hz) | 66.2 ± 5.7 | 40.6 ± 7.3 | $p=0.017$ |

For descriptions of each of the measured intrinsic properties, see the methods section. The n values represent the number of PV+ interneurons. AP frequency was measured from APs evoked by 1 s depolarizing current steps (400 pA). Abbreviations: RMP, resting membrane potential; C<sub>m</sub>, membrane capacitance; R<sub>input</sub>, input resistance.

Next, we tested whether the decreased AP firing frequency of dentate PV+ interneurons from TLE mice continued for months following status epilepticus induction. We examined the firing properties of dentate PV+ interneurons of sham control and TLE mice 4 months after intrahippocampal saline and kainate injection, respectively. Whole-cell patch-clamp recordings revealed that there was a significant decrease in AP frequency of dentate PV+ interneurons from TLE mice (**Fig. 2E, F**; two-way ANOVA with repeated measures; significant IHK treatment main factor, *p*=0.0457; significant IHK treatment×injected current amplitude interaction, *p*<0.0001; Sham controls: n=9 from 6 mice; TLE: n=12 from 5 mice). Since the data from PV+ interneurons were similar between the ipsi- and contra-lateral dentate gyrus (two-way ANOVA with repeated measurements; IHK treatment main factor, *p*=0.608), the results from dentate PV+ interneurons from both hemispheres were combined. The frequency rate of APs induced by 400 pA depolarizing current steps was 40.6±7.3 Hz (n=12) in dentate PV+ interneurons from TLE mice, which was significantly lower than 66.2±5.7 Hz (n=9; *p*=0.017) in sham control mice. In addition, dentate PV+ interneurons from TLE mice showed a numerical decrease in rheobase (TLE: 240.0±25.1 pA, n=12; Sham controls: 191.1±28.1 pA, n=9), which is consistent with the data obtained 4-6 weeks after status epilepticus induction. However, the difference in rheobase between the two cohorts did not reach statistical significance (*p*=0.165). These findings suggest that the reduction in dentate PV+ interneuron firing rate associated with epilepsy development continues in TLE mice for several months following status epilepticus.

### 3.3. Synaptic excitation of dentate PV+ interneurons of TLE mice

Increased synaptic excitation of dentate PV+ interneurons develops in a mouse model of traumatic brain injury along with reduced AP frequency rate (Kang et al., 2022). We thus examined whether synaptic excitation of PV+ interneurons was also upregulated in TLE mice 4-6 weeks after IHK injection. Whole-cell patch-clamp recordings from dentate PV+ interneurons revealed that there were no changes in the frequency (TLE: 12.6±2.1 Hz, n=11 from 5 mice; Sham controls: 9.7±2.3 Hz, n=10 from 5 mice; *p*=0.376) or amplitude (TLE: 36.3±4.7 pA, n=11 from 5 mice; Sham controls: 27.3±2.8 pA, n=10 from 5 mice; *p*=0.124) of sEPSCs in dentate PV+ interneurons (**Fig. 3**). These results suggest that synaptic excitation of dentate PV+ interneurons remains unchanged in TLE mice.

**Figure 3.**
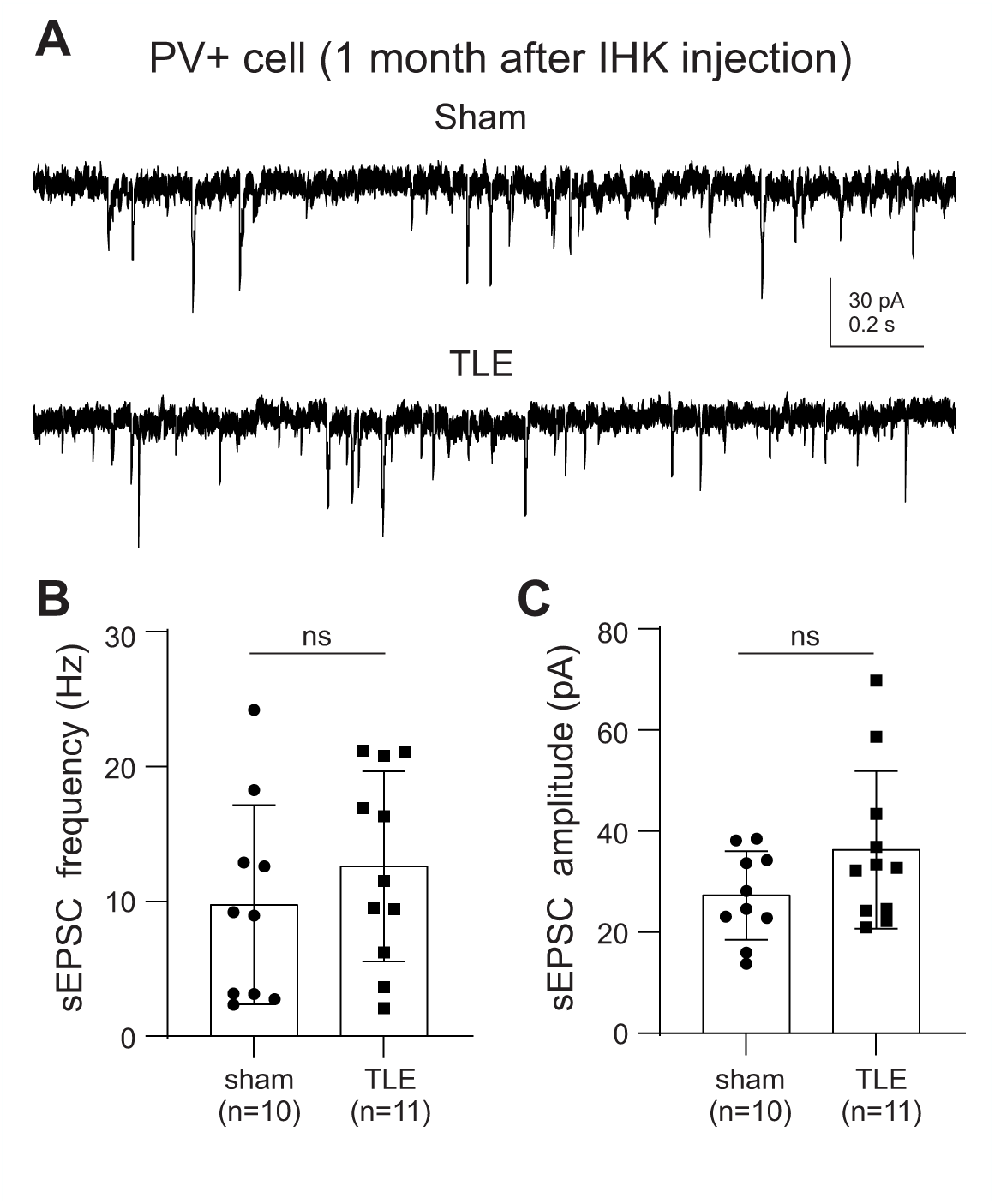
Synaptic excitation of dentate PV+ interneurons. **(A)** Examples of sEPSCs in dentate PV+ interneurons of sham control and TLE mice. **(B, C)** Summary of sEPSC frequency and amplitude. Solid circles and squares represent individual PV+ interneurons. Mean±SEM; ns, not significant.

### 3.4. Synaptic inhibition of DGCs of TLE mice

PVBCs and AACs are major subtypes of GABAergic interneurons in the dentate gyrus (Pelkey et al., 2017). Given that TLE mice showed decreased intrinsic excitability of surviving PV+ interneurons (**Fig. 2**) and loss of PV+ interneurons (**Fig. 1**), we hypothesized that DGCs would receive reduced synaptic inhibition after IHK treatment. We thus sought to determine whether TLE mice manifested reduced AP-dependent GABAergic inhibition in DGCs 4-6 weeks after IHK injection. Whole-cell patch-clamp recordings from DGCs revealed that there was indeed a significant decrease in the frequency of sIPSCs in TLE mice (**Fig. 4A,B**; TLE: 1.95±0.27 Hz, n=28 from 9 mice; Sham controls: 3.86±0.48 Hz, n=25 from 9 mice; *p*=0.0006). However, we found that the amplitude of sIPSCs was significantly greater in TLE mice (**Fig. 4A,C**; TLE: 30.32±1.77 pA, n=28; Sham controls: 22.80±1.47 pA, n=25; *p*=0.004). These results suggest not only that reduced intrinsic excitability and loss of dentate PV+ interneurons of TLE mice contribute to reduced frequency of sIPSCs in DGCs, but also that an increase in sIPSC amplitude develops in TLE mice as a potentially compensatory mechanism.

**Figure 4.**
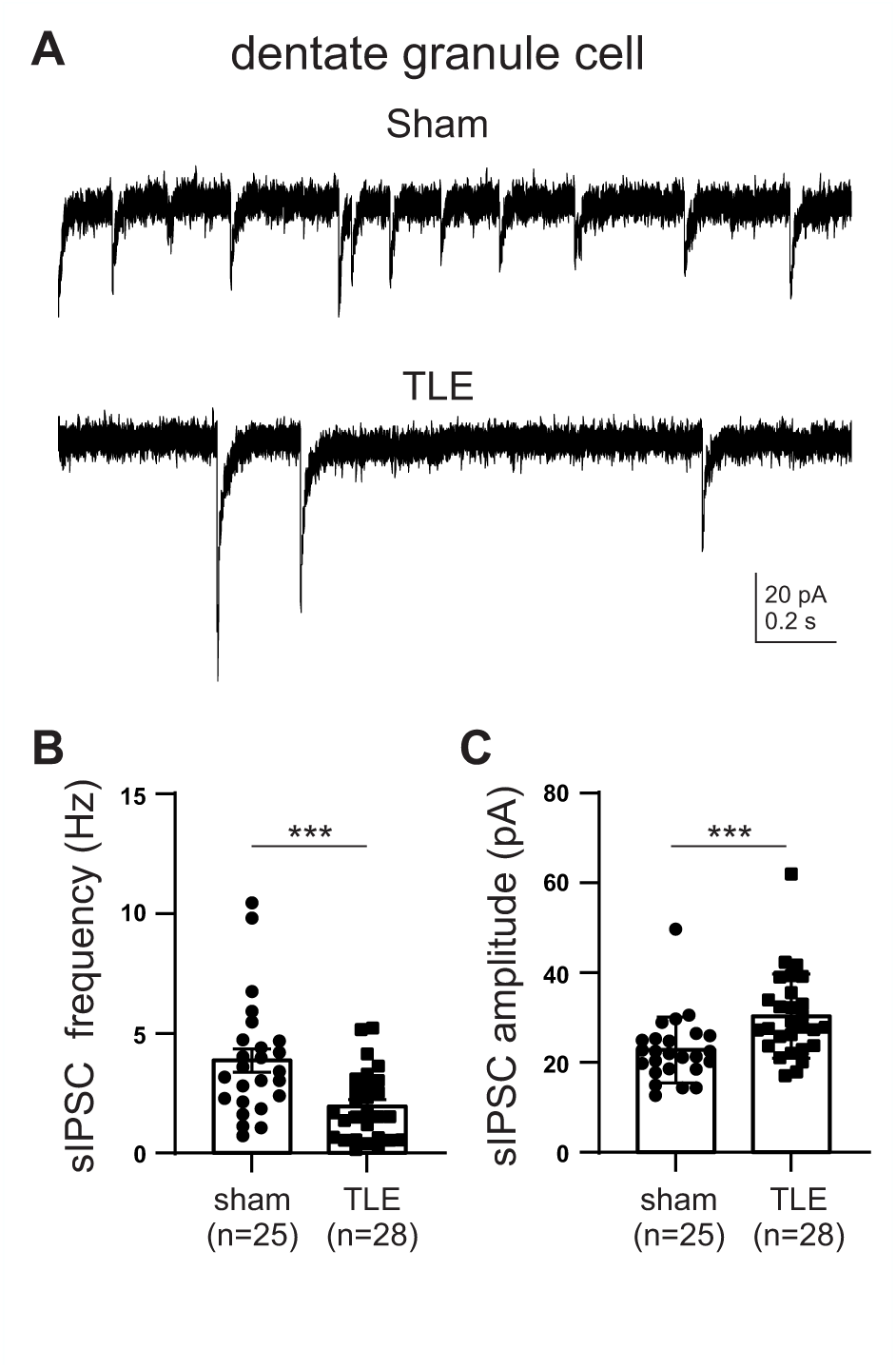
Compromised GABAergic inhibition in DGCs of TLE mice. **(A)** Examples of sIPSCs in DGCs from sham control and TLE mice. Note that there is a decrease in sIPSC frequency in DGCs from TLE mice, whereas there is an increase in sIPSC amplitude in DGCs from TLE mice. **(B, C)** Summary of sIPSC frequency and amplitude. Solid circles and squares represent individual DGCs. Mean±SEM; \*\*\**p*<0.005.

### 3.5. PV+ interneuron mediated inhibition of DGCs from TLE mice

We used PVCre::ChR2 mice to directly measure alterations in PV+ interneuron mediated inhibition in DGCs from TLE mice 4 months after IHK injection. Optogenetic activation of dentate PV+ interneurons from TLE mice was expected to produce fewer APs in PV+ interneurons since there was reduced intrinsic excitability of dentate PV+ interneurons after IHK treatment (**Fig. 2**). Indeed, optogenetic activation of dentate PV+ interneurons of TLE mice by blue light (2 ms duration) produced fewer APs in PV+ interneurons from TLE mice (**Fig. 5A,B**; TLE: 2.5±0.9 APs, n=14 from 9 mice; Sham controls: 3.1±0.2 APs, n=14 from 5 mice; *p*=0.0469). These results are consistent with those from evoked APs using depolarizing current pulses (**Fig. 2**). Next, we investigated whether a decrease in the number of evoked APs in PV+ interneurons resulted in a reduction in PV+ interneuron mediated inhibition of DGCs from TLE mice. Surprisingly, a similar light stimulation caused a greater eIPSC amplitude in DGCs from TLE mice (**Fig. 5C,D**; TLE: 652.4±60.6 pA, n=37 from 6 mice; Sham controls: 536.4±31.7 pA, n=30 from 7 mice; *p*=0.0335). Nonetheless, these results are consistent with the greater amplitude of sIPSCs detected in DGCs from TLE mice (**Fig. 4A,C**).

**Figure 5.**
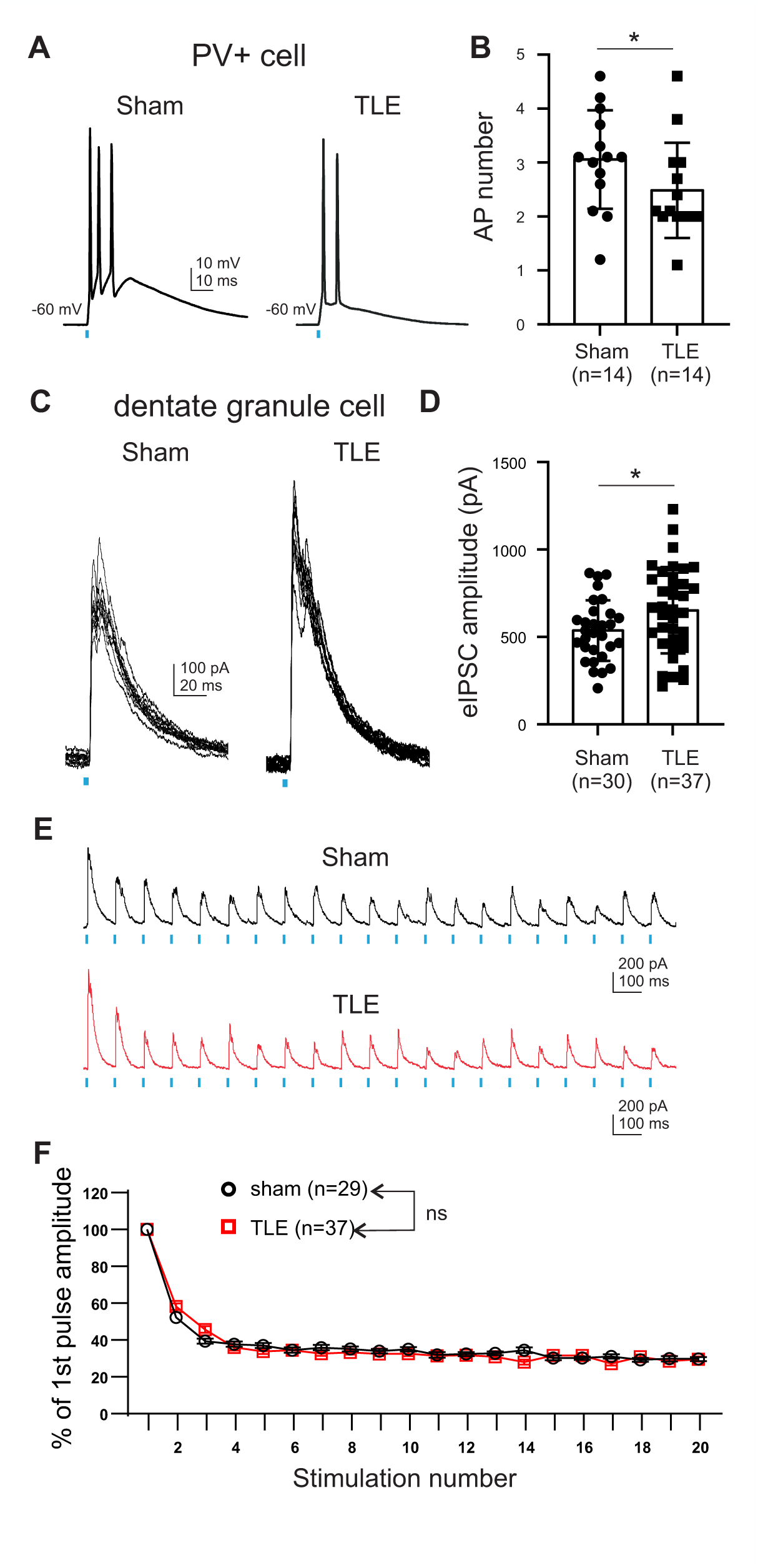
Increased evoked PV+ interneuron mediated IPSCs in DGCs from TLE mice. **(A,B)** Light evoked AP discharges in dentate PV+ interneurons in sham control and TLE mice. Notably, TLE mice elicited lower AP counts in response to comparable light pulses (A). Summary of evoked AP numbers (B). Solid circles and squares represent individual PV+ basket cells. **(C,D)** Representative evoked IPSCs (eIPSCs) in DGCs of sham control and TLE mice (C). Ten eIPSCs were elicited by 0.033 Hz light pulses. Summary of eIPSC amplitude in DGCs (D). Solid circles and squares represent individual DGCs. **(E, F)** Twenty light pulses at a frequency of 10 Hz elicited eIPSCs in DGCs (E). Summary of % of 1st pulse amplitude (F). Mean±SEM; \**p*<0.05; ns, not significant.

Next, we examined whether DGCs manifested altered short-term plasticity of eIPSCs in DGCs from TLE mice using trains of theta-frequency light stimuli (*i.e.*, 20 light pulses at 10 Hz). Given that there is downregulated short-term depression of eIPSCs in DGCs when PV+ interneurons are activated by sustained stimulation in kindled mice (Hansen et al., 2017), it is interesting that such reduced short-term depression of eIPSCs was not observed in DGCs in TLE mice (**Fig. 5E, F**; two-way ANOVA with repeated measures, IHK treatment main factor, *p*=0.6120). These results suggest that the increased PV+ interneuron mediated inhibition of DGCs from TLE mice is not due to epilepsy-associated changes in presynaptic mechanisms.

### 3.6. Chemogenetic activation of PV+ interneurons stops epileptiform activity in the dentate gyrus of TLE mice

To investigate whether activation of hypoactive dentate PV+ interneurons of TLE mice stops or facilitates epileptiform activity in the dentate gyrus, we employed PVCre::Gq-DREADD mice, which allowed remote, chemogenetic activation of PV+ neurons. We first confirmed that chemogenetic activation of PV+ interneurons increased GABAergic inhibition in DGCs in TLE mice 4-6 weeks after IHK injection. Whole-cell patch-clamp recordings revealed that bath application of CNO (5 μM) increased the frequency of sIPSCs in DGCs from TLE PVCre::Gq-DREADD mice (**Fig. 6A,B**; Control: 2.98±0.60 Hz, n=8 from 4 mice; CNO: 8.56±1.76 Hz, n=8 from 4 mice; *p*=0.0054). The amplitude of sIPSCs in DGCs was unaffected by CNO (**Fig. 6A,C**; Control: 25.67±2.66 pA, n=8 from 4 mice; CNO: 25.23±2.25 pA, n=8 from 4 mice; *p*=0.8370). These findings show that in TLE PVCre::Gq-DREADD mice, CNO enhances GABAergic inhibition in DGCs.

**Figure 6.**
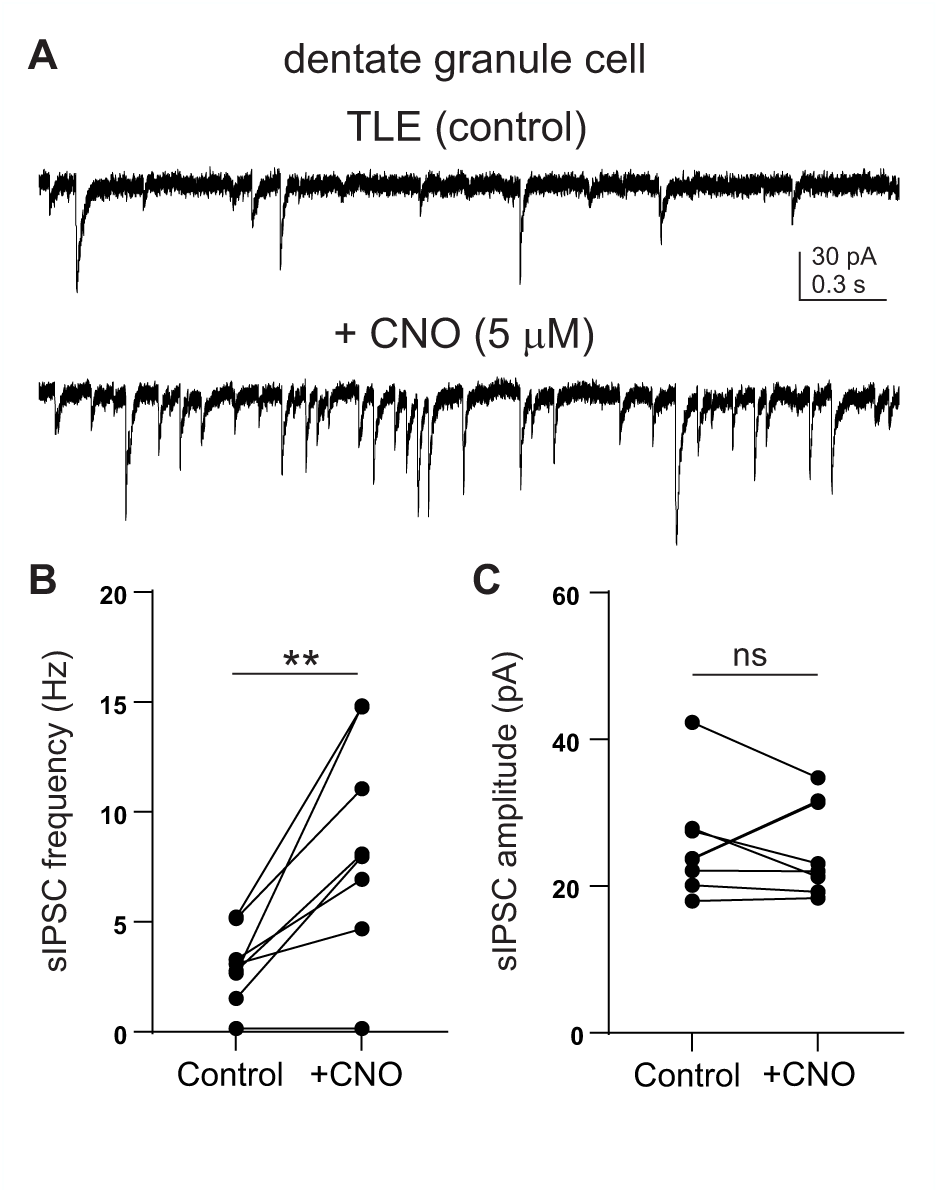
Chemogenetic activation of PV+ interneurons increases sIPSC frequency, but not sIPSC amplitude in DGCs from TLE mice. **(A)** sIPSCs in a DGC of a TLE mouse before and during bath application of CNO (5 μM). CNO application increased sIPSC frequency. **(B, C)** Summary of sIPSC frequency and amplitude. Solid circles represent individual DGCs. Mean±SEM; \*\**p*<0.01; ns, not significant.

Next, we used EEG recordings from TLE PVCre::Gq-DREADD mice with an ipsilateral depth electrode placed in the dentate gyrus to determine whether chemogenetic activation of PV+ interneurons prevented epileptiform activity in the dentate gyrus 4 months after IHK injection. Epileptiform activity was observed in 6 TLE mice prior to CNO treatment (**Fig. 7**). The number of epileptiform activity (**Fig. 7A,B**; Control: 33.7±7.2 per hour, n=6; CNO: 0.7±0.5 per hour, n=6; *p*=0.0313) and the cumulative duration of epileptiform activity (**Fig. 7A,C**; Control: 941.8±238.7s per hour, n=6; CNO: 19.0±13.2s per hour, n=6; p=0.0313) were both decreased by CNO injection (i.p., 1 mg/kg). In contrast, vehicle (i.p., 0.9% saline) had no effects on the number of epileptiform activity (Control: 31.1±3.7 per hour, n=6; Saline: 35.2±4.7 per hour, n=6; *p*=0.0709) or the cumulative duration of epileptiform activity (Control: 731.8±116.1s per hour, n=6; Saline: 849.6±174.7s per hour, n=6; *p*=0.1437) in the same TLE mice. Lastly, in order to determine whether Gq-DREADD expression was necessary for the antiseizure effects of CNO in TLE PVCre::Gq-DREADD mice, we used TLE C57BL/6 mice, who did not express Gq-DREADD. As expected, CNO injection (i.p., 1mg/kg) had no effects on the number of epileptiform activity (Control: 27.7±9.4 per hour, n=4; CNO: 31.4±10.6 per hour, n=4; *p*=0.3960) or the cumulative duration of epileptiform activity (Control: 543.3±230.0s per hour, n=4; CNO: 602.9±250.6s per hour, n=4; *p*=0.2558) in TLE C57BL/6 mice. These findings suggest that chemogenetic activation of PV+ interneurons compensates for reduced GABAergic inhibition, which in turn inhibits epileptiform activity in the dentate gyrus.

**Figure 7.**
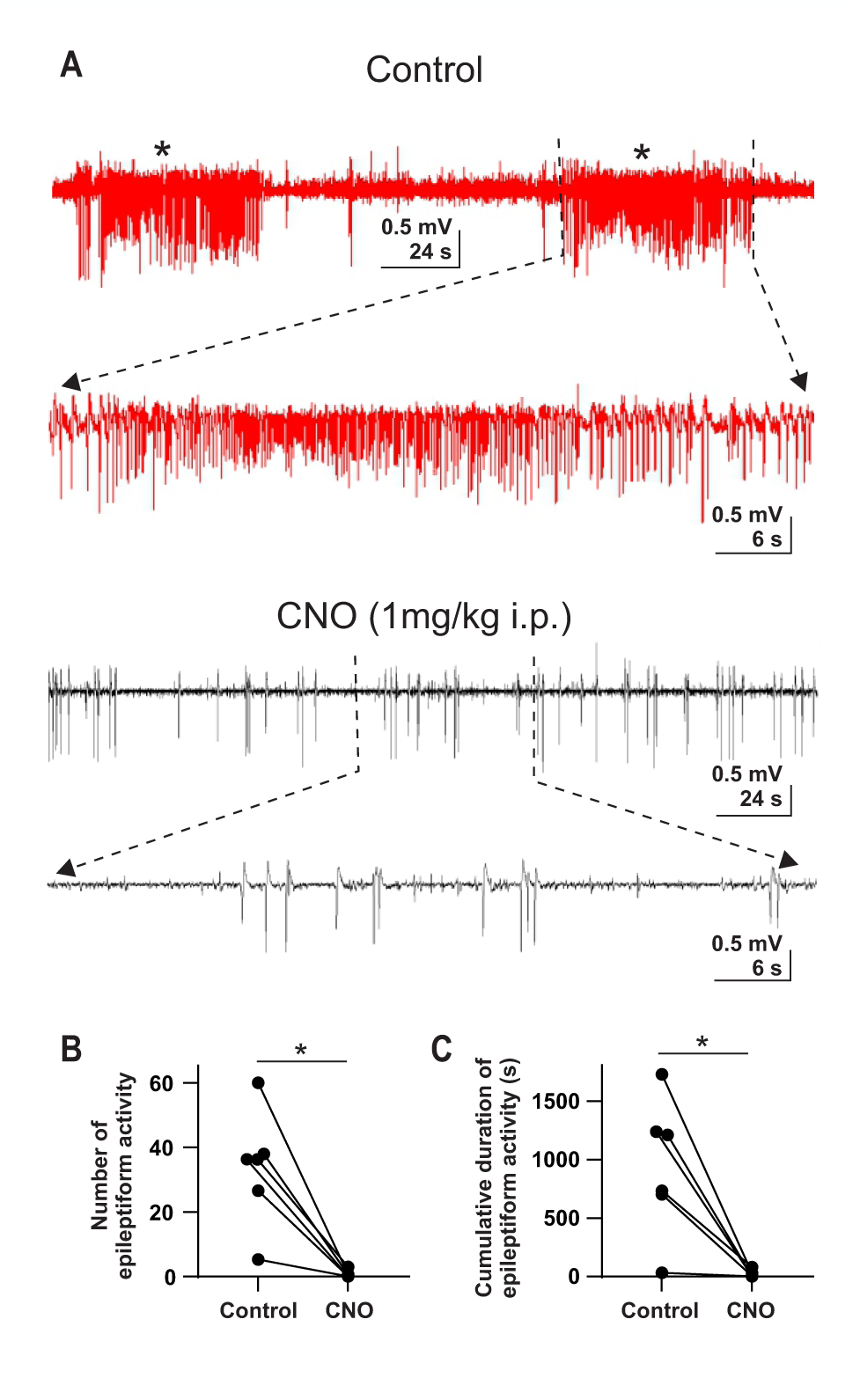
Chemogenetic activation of PV+ interneurons prevents *in vivo* epileptiform activity in the dentate gyrus of TLE PVCre::Gq-DREADD mice. **(A)** Red voltage records show 2 epileptiform activity events (asterisks) before administration of CNO (i.p., 1 mg/kg). Expanded time scale of the epileptiform activity displays high-frequency spikes with high amplitude. The mouse showed only interictal spikes after receiving CNO (black voltage records). Expanded temporal scale of interictal spikes demonstrates low frequency interictal spikes. **(B,C)** Graphs showing significant decreases in the number (B) and cumulative duration (C) of epileptiform activity by CNO treatment (i.p., 1 mg/kg) in TLE PVCre::Gq-DREADD mice (n=6). Epileptiform activity was measured for the 3 hours before CNO administration (control) and the 3 hours after CNO administration. Mean±SEM; \**p*<0.05.

## 4. Discussion

This study showed increased dentate PV+ interneuron mediated inhibition and antiseizure effects of activating hypoactive dentate PV+ interneurons in TLE mice. While our current results are consistent with prior studies showing reduced intrinsic excitability of PV+ interneurons after epileptogenic brain insults (Kang et al., 2022; Khan et al., 2018; Pothmann et al., 2019; Proddutur et al., 2023), novel aspects of PV+ interneuron-associated upregulation of dentate inhibition in TLE and unaltered synaptic excitation and presynaptic release properties of dentate PV+ interneurons are defined. These results suggest that reduced AP generation in dentate PV+ interneurons contributes to hyperexcitability and spontaneous seizures in TLE and normalizing the epilepsy related reduction in AP generation in dentate PV+ interneurons reduces seizures in TLE.

### 4.1. Persistently reduced AP generation in dentate PV+ interneurons, hyperexcitability, and seizures in TLE mice

Reduced intrinsic excitability of dentate PV+ interneurons can impair powerful dentate PV+ interneuron mediated perisomatic inhibition of DGCs in TLE. Importantly, the current study shows that dentate PV+ interneurons manifest a decrease in AP firing rate in the chronic phase of TLE, complementing results obtained shortly after experimental status epilepticus (Proddutur et al., 2023). Therefore, reduced AP generation of dentate PV+ interneurons may play a critical role in epileptogenesis and during the chronic phase of TLE.

Nav1.1 is the primary voltage-gated sodium channel in PV+ interneurons (Ogiwara et al., 2007; Yu et al., 2006) and Nav1.1 immunoreactivity is significantly downregulated in TLE (Qiao et al., 2013). Reduced AP generation of dentate PV+ interneurons may thus be due to downregulated Nav1.1 activity in PV+ interneurons of TLE mice. The functional abnormalities in PV+ interneurons described here (i.e., reduced AP firing rate, increased rheobase, and increased AP threshold) are reminiscent of those commonly found in mouse models of Dravet syndrome (Kaneko et al., 2022; Ogiwara et al., 2007; Rubinstein et al., 2015; Yu et al., 2006). Dravet syndrome is a rare disease, with most patients having a mutation in *SCN1A*, which encodes Nav1.1 (Wu et al., 2015). However, the persistent PV+ interneuron-associated changes in the present TLE model are in sharp contrast to the transient characteristics of impaired AP generation of PV+ interneurons in Dravet syndrome, which are defined by the recovery of fast-spiking features of PV+ interneurons in the chronic phase of Dravet syndrome (Kaneko et al., 2022). Given that correcting the reduced AP generation in PV+ interneurons by genetic or pharmacological approaches targeting Nav1.1 can prevent hyperexcitability and seizures in other neurological disorders (e.g., Dravet syndrome, Alzheimer’s disease) (Richards et al., 2018; Verret et al., 2012; Yamagata et al., 2020), normalizing the reduced AP generation of dentate PV+ interneurons, or utilizing approaches targeting Nav1.1 channel function, may modify epileptogenesis and prevent hyperexcitability and seizures in the chronic phase of TLE.

Other, non-Nav1.1-associated changes in dentate PV+ interneurons could lead to reduced AP firing rate in this interneuron population in TLE. For example, Kv3 K^+^ channels are highly and selectively expressed in the axons of PV+ interneurons and are critical for the characteristic, fast-spiking property of these neurons (Du et al., 1996; Hu et al., 2014). It is thus possible that Kv3 channel-associated changes in PV+ interneurons in TLE mice contribute to the reduced AP firing rate of dentate PV+ interneurons, since loss or downregulation of Kv3 K^+^ channels in PV+ interneurons reduces AP firing rates (Lau et al., 2000; Medrihan et al., 2020). Moreover, recent studies showed that a Kv3 positive modulator reversed reduced AP generation of Kv3.1-expressing neurons and reduced seizures in a mouse model of a progressive myoclonus epilepsy (Feng et al., 2024), suggesting that enhancing Kv3 activity in PV+ interneurons could be a novel therapeutic target. Importantly, human epilepsy has been associated with rare variations found in the *KCNC* genes that encode Kv3 channels (Li et al., 2021; Muona et al., 2015). Therefore, it will be crucial to determine in future studies whether and how functional alterations in Nav1.1 and Kv3 channel function result in decreased intrinsic excitability of dentate PV+ interneurons in TLE.

### 4.2. Increased PV+ interneuron mediated inhibition in TLE

This study shows evoked PV+ interneuron mediated inhibition in the dentate gyrus is increased in TLE mice. These results agree with prior evidence showing not only that PV+ interneurons in the dentate gyrus experience an increase in symmetrical synapses on axon initial segments of DGCs, maintenance of a similar number of synapses on DGC somata, and axon sprouting in TLE (Christenson Wick et al., 2017; Thind et al., 2010; Wittner et al., 2001), but also that dentate PV+ interneuron-associated feedback inhibition is upregulated in a mouse model of posttraumatic epilepsy (PTE) after activation of DGCs (Kang et al., 2022). Increased evoked dentate PV+ interneuron mediated inhibition has an important implication for treatments of spontaneous seizures in TLE since the dentate gyrus is a prominent location of pathology in TLE.

Short-term plasticity of PV+ interneuron mediated eIPSCs in DGCs was unaltered after IHK treatment, which agrees with prior studies in the dentate gyrus following epileptogenic insults (Hansen et al., 2017; Proddutur et al., 2023). These results suggest that the increased PV+ interneuron mediated inhibition of DGCs in IHK mice is mainly driven by epilepsy-associated changes in postsynaptic sites of PV+ interneurons rather than presynaptic GABA release onto DGCs, and axon sprouting-associated increase in GABAergic synapses may also contribute.

Indeed, experimental models of TLE show axon sprouting of PV+ and other interneurons, increased GABAergic synapses, increased number of GABA_A_ receptors at inhibitory synapses on somata and axon initial segments, increased amplitude of sIPSCs, and increased tonic inhibition of DGCs (Boychuk et al., 2022; Cossart et al., 2001; Nusser et al., 1998; Thind et al., 2010). Our current findings of increased PV+ interneuron mediated evoked inhibition are consistent with these previous results. The underlying molecular and cellular mechanisms of increased PV+ interneuron mediated evoked inhibition of DGCs in TLE mice will need to be identified in future studies.

### 4.3. Functional relevance

Increasing intrinsic excitability of hypoactive dentate PV+ interneurons may be a novel target of seizure treatments in acquired epilepsies including TLE and PTE. The "dormant dentate basket cell" theory (Sloviter, 1987; Sloviter, 1991) is not supported by our main findings of reduced AP generation of dentate PV+ interneurons in TLE, as this study did not suggest a decrease in the frequency and amplitude of sEPSCs in dentate PV+ interneurons in the TLE mice. However, dentate PV+ interneurons may become hypoactive in TLE as a result of several epilepsy-associated activity changes (e.g., decreased intrinsic excitability). PV+ interneurons critically participate in feedback and feedforward inhibition to prevent hippocampal hyperexcitation (Lee et al., 2019). Reduced intrinsic excitability of dentate PV+ interneurons in TLE may thus lead to reduced feedback and feedforward inhibition, thus contributing to dentate gyrus hyperexcitability and spontaneous seizures in TLE. Indeed, reduced feedback inhibition in the CA1 region and feedforward activation of dentate PV+ interneurons have been implicated in hyperexcitability and increased seizure propagation in TLE and traumatic brain injury (Folweiler et al., 2020; Pothmann et al., 2019). The present studies show that *in vivo* chemogenetic activation of PV+ interneurons significantly reduces TLE-associated epileptiform discharges in the dentate gyrus. These results disagree with prior findings showing that PV+ interneurons promote hippocampal epileptiform activity and paradoxical rebound excitation in other seizure models (Ellender et al., 2014; Wang et al., 2022). However, they agree with recent studies showing that opto- or chemogenetic activation of PV+ interneurons produces antiseizure effects in TLE (Krook-Magnuson et al., 2013; Wang et al., 2018).

In conclusion, the current findings demonstrate that while stimulation of dentate PV+ interneurons produces enhanced inhibition of DGCs in the chronic phase of TLE, they manifest a long-term reduction in intrinsic excitability that suppresses their ability to inhibit DGC activity. It is unclear whether these altered circuit dynamics represent compensatory changes to address reduced overall synaptic inhibition in TLE or if they contribute to seizure susceptibility and behavioral comorbidities of TLE. This study highlights the persistent reduction of intrinsic excitability of dentate PV+ interneurons that may present future opportunities for developing novel treatments for TLE by targeting molecules selectively expressed in GABAergic interneurons (e.g., Nav1.1 and Kv3 K^+^ channels), but not excitatory cells to achieve cell type-specific pharmacological modulation.

## Acknowledgments

This work was supported by R01 NS092552 (to B.N.S) and CVMBS Shared Research Program at Colorado State University (to S.H.L.). We thank Dr. Seonil Kim for generous help with the tile-scanned imaging system, Dr. Katalin Smith for help with tissue processing, and Evelina Bouckova for help with behavioral seizure analyses. The sponsors had no role in study design, data collection, analysis and interpretation, or writing of this manuscript.

## Author contributions

S.H.L., Y.J.K., and B.N.S. designed research; S.H.L. and Y.J.K. performed research and analyzed data; S.H.L., Y.J.K., and B.N.S. wrote the paper.

## Abbreviations

AACs: axo-axonic cells
AP: action potential
BIO: biocytin
Cm: membrane capacitance
CNO: clozapine N-oxide
DGCs: dentate granule cells
eIPSCs: evoked inhibitory postsynaptic currents
EYFP: enhanced yellow fluorescent protein
GCL: granule cell layer
H: hilus
IHK: intrahippocampal kainite
ML: molecular layer
PTE: posttraumatic epilepsy
PVBCs: PV+ basket cells
PV+: parvalbumin-positive
R_input_: input resistance
RMP: resting membrane potential
sEPSCs: spontaneous excitatory postsynaptic currents
sIPSCs: spontaneous inhibitory postsynaptic currents
TLE: temporal lobe epilepsy
TOM: tdTomato.

